# Biphasic Changes in Hippocampal Granule Cells after Traumatic Brain Injury

**DOI:** 10.1101/2024.01.23.576807

**Authors:** Joanna Danielewicz, Nerea Llamosas, Irene Durá, Danillo Barros de Souza, Serafim Rodrigues, Juan Manuel Encinas-Pérez, Diego Martin Mateos

## Abstract

Traumatic brain injury (TBI) leads to a wide range of long-lasting physical and cognitive impairments. Changes in neuronal excitability and synaptic functions in the hippocampus have been proposed to underlie cognitive alterations. The dentate gyrus (DG) acts as a “gatekeeper” of hippocampal information processing and as a filter of excessive or aberrant input activity. In this study, we investigated the effects of controlled cortical impact, a model of TBI, on the excitability of granule cells (GCs) and excitatory postsynaptic transmission in the DG at three time points, 3 days, 15 days and 4 months after the injury. Our results indicate that changes in the short term are related to intrinsic properties, while changes in the long term are more related to input and synaptic activity, in agreement with the notion that TBI-related pathology courses with an acute phase and a later long-term secondary phase. A biphasic response, a reduction in the shorter term and an increase in the long term, was found in TBI neurons in the frequency of sEPSC. These changes correlated with a loss of complexity in the pattern of the synaptic input, an alteration that could therefore play a role in the chronic and recurrent TBI-asssociated hyperexcitation.

## 1. Introduction

Traumatic brain injury (TBI) is a major cause of morbidity and mortality worldwide, affecting millions annually. TBI leads to diverse histopathological and behavioral consequences that begin shortly after trauma and can persist as chronic symptoms months or years later. Following the initial insult, direct mechanical damage to brain tissue triggers an acute phase characterized by neuroinflammation, oxidative stress, excitotoxicity, and gliosis ^1–3^. The acute phase is followed by a secondary phase where some effects persist subtly and new issues arise. Neuronal hyperexcitation, due to underlying changes in circuits, is common in this phase and can lead to epileptiform activity or even seizures ^4^. It is also associated with TBI-related neuropsychological symptoms such as memory loss, learning deficits, and alterations in sensorimotor function and personality ^5–7^.

Synaptic disruption occurs both short- and long-term post-TBI, but the mechanisms driving long-term effects are not well understood. The hippocampus is particularly vulnerable to TBI due to its recurrent circuits, and its disruption plays a role in impairments of learning, memory, and higher cognitive functions ^8^. Similar chronic neuropathological changes occur in humans and experimental animal models of TBI ^9–11^. Within the hippocampus, the dentate gyrus (DG) acts as a “gatekeeper” filtering excessive or aberrant input activity through three main mechanisms: (1) intrinsic membrane properties of granule cells (GCs) that resist excessive excitation; (2) interconnections between GCs that prevent synchronization; and (3) surrounding inhibitory interneurons and synaptic inhibitory efficacy ^12–14^. Thus, GCs in the hippocampus play a pivotal role in the consequences of TBI.

Studies on DG electrophysiology have consistently shown that TBI increases DG excitability in rodents, reducing its filtering capacity. DG hyperexcitability may result from increased excitability of surviving granule cells, decreased synaptic inhibitory efficacy, and/or increased synaptic excitatory efficacy ^15–17^. This study aims to determine the effects of TBI on granule cell excitability and excitatory synaptic activity in the DG at short (3 days), medium (15 days), and long (4 months) terms. We used various analytical tools, such as principal component analysis (PCA) and Lempel-Ziv complexity (LZC), to gain further insight. Importantly, to better isolate the effects of brain tissue damage, we included a pure control group without craniotomy in addition to the standard sham group.

## 2. Materials and Methods

### Animals and surgery

8 weeks old C57BL/6 male and female mice were used for all procedures. Mice were housed at constant humidity and temperature with a 12-h light/dark cycle with food ad libitum. All experimental procedures were approved by the University of the Basque Country (EHU/UPV) Ethics Committee (Leioa, Spain) and the Comunidad Foral de Bizkaia (CEEA: M20/2015/236).

The animals were randomly assigned to the TBI, sham or control group. TBI mice model was established with a controlled cortical impact (CCI) method (TBI 0310, Precision Systems and Instrumentation). Briefly, mice were anesthetized with ketamine and fixed on stereotaxic apparatus. A midline incision was made to expose the skull. An approximate 4-mm craniotomy was performed over the left parietotemporal cortex using a motorized drill. The CCI was centered at 2.5-mm lateral to the midline and at 0.5-mm anterior to bregma and was conducted with a 30-mm flat-tip impounder (velocity-5.5 m/s; duration-600 ms; depth-1 mm). After CCI, the scalp incision was sutured, and the mice were placed on a heating pad until anesthesia recovery. Sham mice received all these procedures, except for CCI. Control group of animals was subjected only to ketamine anesthesia (n=21 mice for control group; n=16 mice for sham group and n= 21 mice for TBI group).

### Preparation of brain slices

Acute coronal hippocampal slices (250 μm) were cut using standard methods as previously described ^18^. Briefly, mice were decapitated, and brains rapidly removed in ice-cold cutting solution containing (in mM): 92 NMDG, 2.5 KCl, 0.5 CaCl_2_, 10 MgSO_4_, 1.25 NaH_2_PO_4_, 30 NaHCO_3_, 20 HEPES, 5 Na_-_ascorbate, 3 Na-pyruvate, 2 Thiourea and 25 D-glucose (pH 7.3 – 7.4; 300 – 310 mOsm) and bubbled with the mixture of 95% O2– 5% CO2. Slices were stored submerged at room temperature for recovery.

### Whole-cell recordings

After 1-1.5 h individual slices were placed in the recording chamber mounted on the stage of Scientifica microscope and superfused at 3 ml/min with warm (32 ± 0.5 °C), modified ACSF of the following composition (in mM): 124 NaCl, 4.5 KCl, 1.25 NaH_2_PO_4_, 26 NaHCO_3_, 1 MgSO_4_ * 7 H_2_O, 1.8 CaCl_2_, and 10 D-glucose (pH 7.3-7.4; 300-310 mOsm), bubbled with the mixture of 95% O2–5% CO2. The pipette solution contained (in mM): 125 K-gluconate, 20 KCl, 2 MgCl2, 10 HEPES, 4 Na2-ATP, 0.4 Na-GTP, 5 EGTA (pH 7.3-7.4; 295-305 mOsm,). Pipette resistance was 7-9 MΩ. The calculated liquid junction potential using this solution was 13.1 mV, and data were corrected for this offset. Signals were recorded using Axon MultiClamp 700B amplifier (Molecular Devices), filtered at 2 kHz, and digitized at 20 kHz using Digidata 1550A (Molecular Devices) interface and Clampex 10 software (Molecular Devices, USA).

### Lempel-Ziv complexity

To analyze changes in the dynamics of spontaneous excitatory postsynaptic currents(sEPSCs) sequences across different groups and time scales, we used a non-linear measure of complexity known as Lempel-Ziv complexity (LZC) ^19^. Each sEPSC sequence was binarized using the median of the signal as the threshold. LZC was then calculated on the binary sequence (for detailed LZC calculation see the supplementary material). We compared the LZC values between states and over time. Finally, we examined the correlation between the LZC values and the mean sEPSC.

### Principal component analysis

We applied principal component analysis (PCA) ^20^ to the set of properties obtained from the passive and intrinsic membrane properties. This analysis allowed us to reduce the dimensionality problem to two principal components, facilitating comparisons between groups and the three different time points (see supplementary material for details).

### Statistical analysis

Statistical analysis was performed using GraphPad Prism 9 (GraphPad Software) or Phyton (Python Software Foundation). All data sets were tested for deviation from normal distribution (D’Agostino). Data were analyzed using either a one – way ANOVA or two – way ANOVA. Post – hoc analysis for ANOVA was conducted using Tukey’s test. For data that were not normally distributed, nonparametrical Kruskal-Wallis test was performed followed by Dunńs multiple comparison test. Cumulative probability distribution was compared using Kolmogorov–Smirnov’s test. The level of statistical significance was set at p < 0.05.

## 3. Results

We first studied the passive membrane properties of DG granule cells at 3 different timepoints (3 days, 15 days and 4 months) after the surgery, in sham and TBI mice, with the addition of a pure control group that did not have craniotomy. Basal excitability was not changed over time in any group, as resting membrane potential (RMP) (Fig. 1A), membrane constant tau (Fig. 1B) and membrane input resistance (Fig. 1C) values in all groups remained similar in all time points. Next, we sought to determine whether TBI affects the intrinsic excitability of granule cells. Somatic current injections (Fig. 1D) showed that, in general, the sham group fired fewer action potentials (AP) per current step at all time points (Fig. 1E-G).

**Figure 1.**
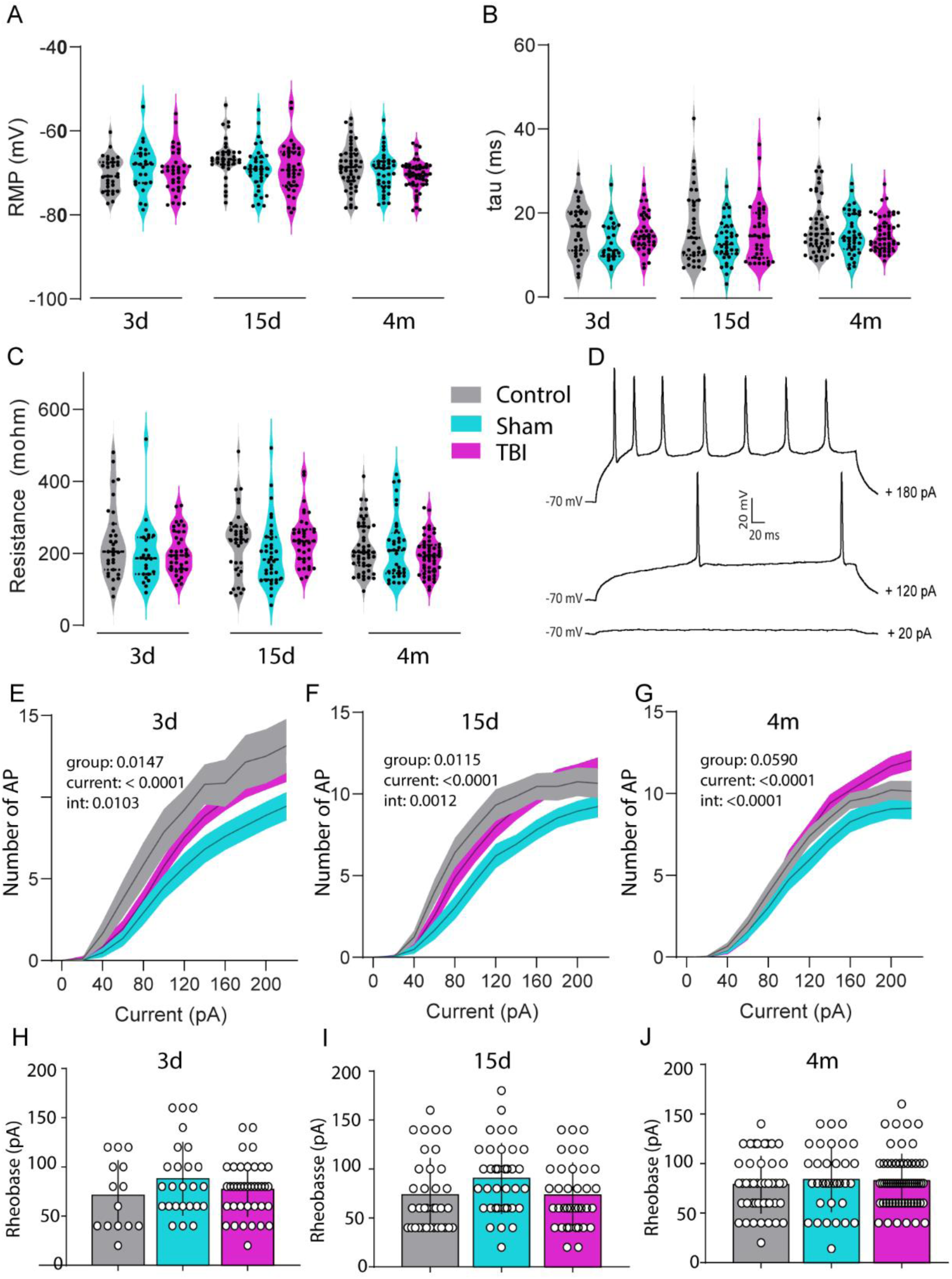
TBI does not alter membrane excitability in granule cells. (**A**) Resting membrane potential of control, sham and TBI granule cells, 3 days, 15 days and 4 months after the surgery. (**B**) *Tau* membrane constant of control, sham and TBI granule cells. (**C**) Input resistance of granule cells from control, sham and TBI groups. (**D**) Representative trace for membrane potential responses to current pulses from +20pA to +220 pA in steps of 20 pA (**E-G**) Input/output curves for control, sham and TBI granule cell population at day 3 (E), day 15 (F) and 4 months (G) after the surgery (Two-way-ANOVA, group F (2, 74)= 4.472, p=0.0147, int F (22, 814)= 1.850, p=0.0103 for day 3; group F (2, 104)= 4.663, p= 0.0115, int F (22, 1144)= 2.192, p= 0,0012 for day 15; group F (2, 138)= 2.890, p= 0.0590, F (22, 1518)= 3.200, p< 0.0001 for 4 months).(**H-J**) Bar plots show rheobase from control, sham and TBI groups at day 3 (H), day 15 (I) and 4 months (J) after the surgery. For A, B and C n=31 neurons from control; 24 from sham and 37 from TBI at day 3; n=36 neurons from control; 36 from sham and 37 from TBI at day 15; n=49 neurons from control; 36 from sham and 52 from TBI at 4 months. For E to J n=14 neurons from control; 25 from sham and 38 from TBI at day 3; n=35 neurons from control; 35 from sham and 37 from TBI at day 15; n=43 neurons from control; 34 from sham and 64 from TBI at 4 months. In violin plots dots represent individual cell values and dashed lines quartiles and medians. For line graphs area fill represents SEM. Circles in bar graphs represent cell values and bars represent mean ± SEM.

To investigate membrane and AP properties with deeper insight, we performed a detailed analysis of RMP, membrane resistance, and evoked AP waveforms. We calculated a total of fifteen variables (Table 1), including amplitude, duration and velocity, from different fractions of the AP elicited by current injections in the three groups at the three time points. Due to the high dimensionality of the data, we chose to use Principal Component Analysis (PCA) as a dimensionality reduction method (Fig. Supplementary 1A-C). We found a significant decrease in the variance of Principal Component 1 (PC1) in TBI neurons compared to control and sham ones at days 3 and 15. Principal Component 2 (PC2) showed that TBI scores were significantly higher than control and sham only at day 15 (Supplementary Fig. 1A-D). Significant effects were found for several variables at different time points, especially for the TBI group, although sham neurons also presented differences versus the control group. At the 3-day time point, TBI had higher AP overshot (Fig. Supplementary 1H) and time to first AP (Fig. Supplementary 1I). The 15-day time point was particularly subject of changes. The TBI group had higher AP half_-_width (Fig. Supplementary 1F), amplitude (Fig. Supplementary 1G), overshot (Fig. Supplementary 1H) plus higher AHP amplitude (Fig. Supplementary 1M). At the 4_-_month time point, TBI AHP duration (Fig. Supplementary 1N) and decay (Fig. Supplementary 1O) was lower. Interestingly, although for the most part the sham group measurements were similar to those of the control group, at particular time points several features changed in alignment with TBI (Fig. Supplementary 1G, H, M, Q) while in occasions was different to both the control and the TBI group (Fig. Supplementary 1I, K, O).

**Table 1.** Variables included in the PCA analysis.

|  |  |
| --- | --- |
| Resting membrane potential (mV) | AP amplitude (mV) |
| Action potential (AP) overshoot (mV) | Membrane resistance (MΩ) |
| AP After hyperpolarization (AHP) amplitude (mV) | Membrane time constant Tau (ms) |
| AP half-width (ms) | AP AHP duration (ms) |
| AP max rise slope (mV/ms) | Voltage displacement $\Delta V_t$ (mv) |
| AP max decay slope (mV/ms) | AP AHP decay (mV*ms) |
| AP threshold (mV) | Time to first AP (ms) |
| Inter spike interval (ms) |  |

We next sought to explore the impact of TBI on the spontaneous excitatory synaptic activity of granule cells (Fig. 2A).

**Figure 2.**
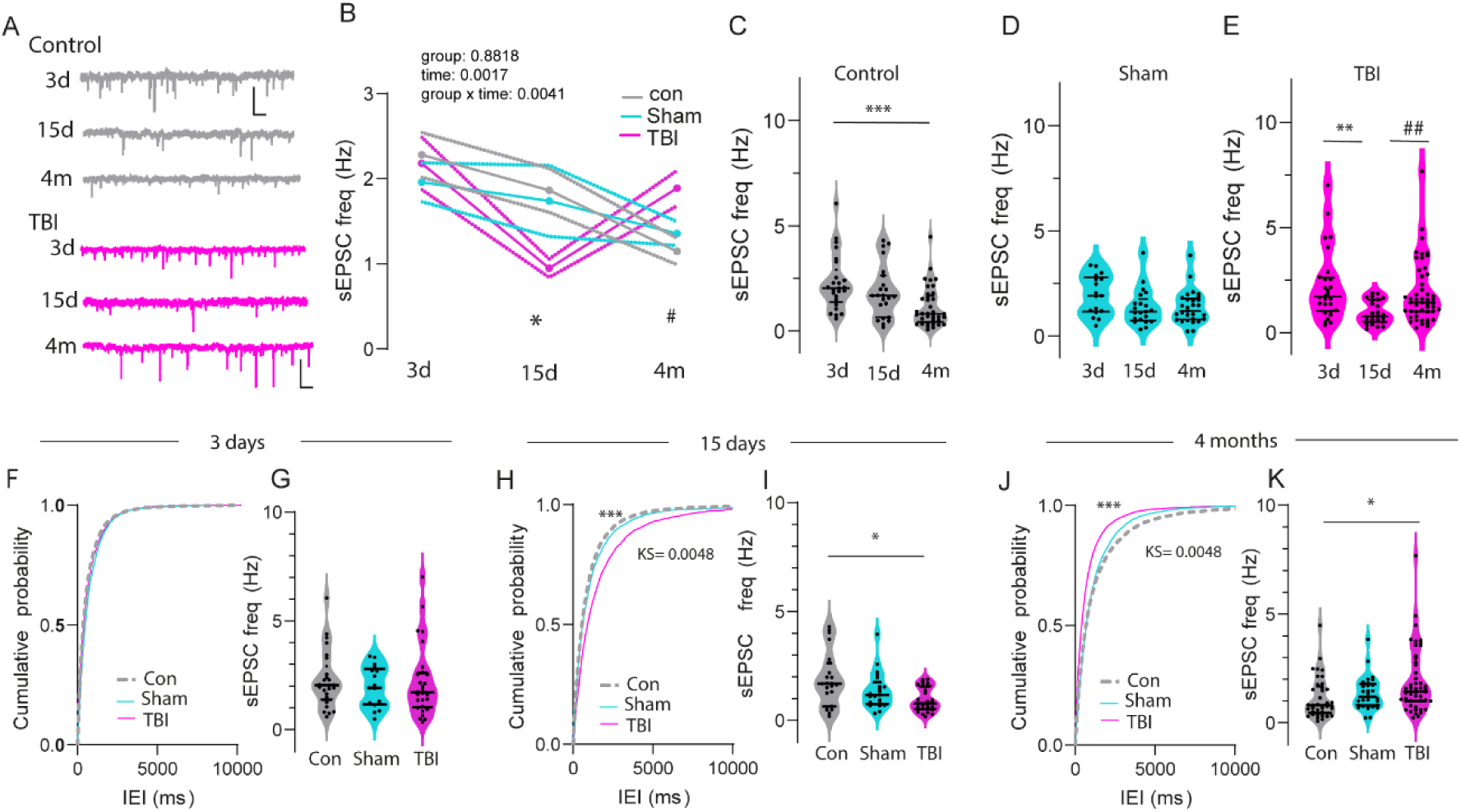
TBI alters the time course of spontaneous synaptic activity and exerts a biphasic modulation of synaptic events. (**A**) Representative sEPSC traces from control and TBI granule cells at day 3, day 15 and 4 months after surgery (Scale bar 20 pA, 500ms). (**B**) sEPSC frequency at the short_-_medium- and long-term for control, sham and TBI granule cells (Two-way-ANOVA, group F (2, 247) = 0.1259, p = 0.8818; int F (4, 247) = 3.935, p= 0.0041; time F(2, 247) = 6,522, p= 0.00171. *p= 0.0334 for TBI vs control and ^#^p=0.0216 for TBI vs control in Tukey’s multiple comparisons test). (**C**) sEPSC frequency in control cells at day 3 (n=25), day 15 (n=23) and 4 months (n=37) after surgery (Kruskal-Wallis test, H(2)= 14.67, p=0.0007. ***p=0.0005 for month 4 vs day 3 in Dunn’s multiple comparisons test). (**D**) sEPSC frequency in s cells at day 3 (n=17), day 15 (n=21) and 4 months (n=29) after surgery. (**E**) sEPSC frequency in TBI cells at day 3 (n=28), day 15 (n=26) and 4 months (n=49) after surgery (Kruskal-Wallis test, H(2)= 13.87, p=0.0010. **p=0.0013 for day 15 vs day 3 and ^##^p=0.0075 for 4 months vs 15 days in Dunn’s multiple comparisons test). For line graphs dots represent group means and area fill represents SEM. Dots in violin plots represent cell median values and dashed lines quartiles and group medians. (**F-G**) Cumulative probability plots of inter_-_event interval (IEI) (F) and violin plots showing sEPSC frequency (G) for control, sham and TBI granule cells at day 3 after the surgery (n=25 for control, n= 17 for sham and n= 28 for TBI). (**H-I**) Cumulative probability plots of IEI (H) and sEPSC frequency violin plots (I) for control, sham and TBI granule cells at day 15 after the surgery. (n=23 for control, n= 22 for sham and n= 26 for TBI. Kruskal-Wallis test, H(2)= 7.792, p=0.0203. *p= 0.0163 for TBI vs control in Dunn’s multiple comparisons test). (**J-K**) Cumulative probability plots of IEI (J) and sEPSC frequency violin plots (K) for control, sham and TBI granule cells 4 months after the surgery (n=37 for control, n= 29 for sham and n= 49 for TBI. Kruskal-Wallis test, H(2)= 8.164, p=0.017. *p= 0.013 for TBI vs control in Dunn’s multiple comparisons test). Dots in violin plots represent cell median values and dashed lines quartiles and group medians.

Intragroup comparisons revealed that in control cells, spontaneous excitatory postsynaptic currents (sEPSC) frequency showed a significant decrease after 4 months attributable to the physiological course of excitatory transmission over the time (Fig. 2B, C). Noticeably, the frequency of sEPSC in the TBI group underwent a clear biphasic change. First, there was a substantial reduction (day 3 to 15) and then a marked increase in the long term (day 15 to 4 months) (Fig. 2, B, E). sEPSC frequency remained stable over the time in sham group (Fig. 2B, D), indicating that the natural decrease observed in controls was counterbalanced by the procedure.

We next performed intergroup comparisons of sEPSC frequency, in terms of mean inter_-_event interval (IEI) per cell and cumulative frequency (the cumulative probability distribution of IEI). No changes were found at the 3-day time point (Fig. 2F, G). However, at day 15, sEPSC frequency of TBI neurons was statistically decreased compared to control neurons (Fig. 2H, I). In contrast, TBI caused a marked increase in sEPSC frequency 4 months after the surgery (Fig. 2J, K). Indeed, an increase in the frequency of sEPSC of granule cells after brain injury compared to uninjured controls has been previously described, but at the earlier timepoint of 8-12 weeks ^21^. Because the amplitude and cumulative probability of the amplitude of sEPSC remained unaltered by time and/or surgery (Supplementary Fig. 2A-G), these results point out to a potential biphasic remodeling of synaptic inputs in granule cells after TBI.

To also test changes in the pattern of synaptic dynamics, we measured the regularity of spontaneous postsynaptic currents by applying Lempel-Ziv complexity (LZC) to sEPSC sequences. Complexity increased naturally over time in the control group, and also in the sham group (Fig. 3A). The complexity in TBI also increased in the 15-day time point, but decreased in the 4-month timepoint showing again a biphasic response (Fig. 3A). In fact, there was a strong negative correlation between sEPSC frequency and complexity in all groups at all time points (Fig.3B) supporting the validity and usefulness of this measurement.

**Figure 3.**
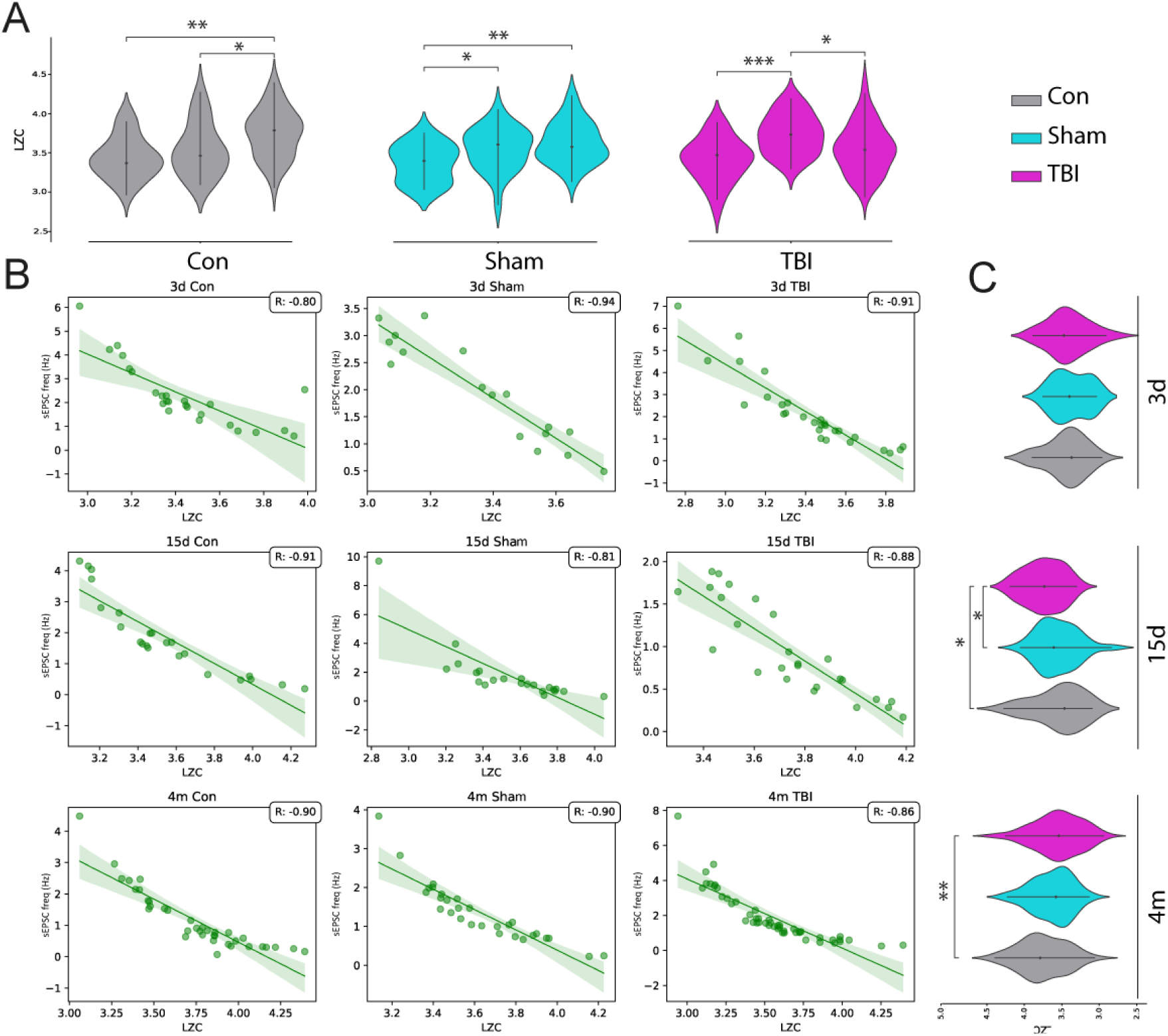
Lempel-Ziv complexity (LZC) analysis of sEPSC sequences. **(A)** Comparison of LZC across different time points. (Kruskal-Wallis test, H(2)= 6.699, p=0.00025. *p= 0.00026 for 3 days vs 4 month in control group; H(2)= 4.325, p=0.0213, *p= 0.0204 for 15 days vs 4 month in control group; H(2)= 5.756, p=0.00052, *p= 0.0059 for 3 days vs 4 month in sham group; H(2)= 4.824, p=0.02721, *p= 0.0281 3 days vs 15 days in sham group; H(2)= 5.324, p=0.00018, *p= 0.00026 for 3 days vs 15 days in TBI group H(2)= 6.654, p=0.0142, *p= 0.0122 for 3 days vs 4 month in TBI group in Dunn’s multiple comparisons test. **(B)** Correlation between LZC and median sEPSC over time and across groups. The correlation value (R) is indicated in the top right corner of each subplot. **(C)** Comparison of LZC between groups at specific time points. (Kruskal-Wallis test, H(2)= 5.793, p=0.0232, *p= 0.0253 for sham vs TBI in 15 days; H(2)= 6.343, p=0.0215, *p= 0.0223 for sham vs TBI in 4month; H(2)= 5.673, p=0.0069, *p= 0.0072 for control vs TBI in 4month in Dunn’s multiple comparisons test). n=26 neurons from control; 18 from sham and 27 from TBI at day 3; n=23 neurons from control; 22 from sham and 25 from TBI at day 15; n=36 neurons from control; 29 from sham and 48 from TBI at 4 months. Violin plots represent group medians and lines SD. In line graphs area fill represents SEM and dots represent cell values.

The intergroup comparisons showed that TBI granule cells, compared to controls, had less complex sEPSC signals 15 days and 4 months after the surgery (Fig.3C). These results could therefore reflect an increase in the number of afferents delivering a more patterned (unvarying) presynaptic glutamate release to granule cells after TBI in the long term. As an internal control, we truncated all sequences longer than 200 points and re_-_analyzed them using the previously described LZC method. Although the absolute values differed, the data remained consistent with the no length-normalized data (Fig. Supplementary 3-C).

Finally, we calculated the correlation between sEPSC frequency and resting membrane potential and found no significant correlation in any group at any time point (Fig. Supplementary 4).

## 4. Discussion

Our study suggests that neither traumatic brain injury (TBI) nor sham surgery may have long-term effects on the passive membrane properties and action potential (AP) generation thresholds of GCs in the dentate gyrus. The stability of resting membrane potential, membrane constant tau, and membrane input resistance over time in all groups supports this notion. Instead, our results support the view that TBI induces a variety of subtle changes that vary temporally rather than producing strong and consistent over-time effects. Notably, the sham surgery group also displayed temporal changes, indicating that it cannot be considered as a pure control group, highlighting the need for careful interpretation of results when using sham surgeries as controls.

Previous research has shown that the early post-traumatic period is crucial in determining outcomes in head trauma ^22^. Animal studies often focus on short-term effects to prevent TBI-induced changes by intervening early. However, electrophysiological, anatomical, and behavioural consequences of TBI can develop over long periods, even up to several months post-injury.

Contrary to commonly reported outcomes, our findings suggest that TBI does not affect the excitability of granule cells in response to current pulses, nor does it alter the rheobase, supporting the notion that different TBI models and electrophysiological approaches can yield varying results ^16,17,23^. Our voltage-clamp recordings of spontaneous current activity reveal significant TBI-induced changes in DG electrophysiology. Age_-_related declines in sEPSC frequency observed in control and sham groups contrast with the biphasic changes in the TBI group, which exhibit an initial decrease followed by a significant increase. These findings align with studies reporting varied effects of TBI on synaptic transmission over time ^24–26^. In the chronic period, increased spontaneous synaptic excitation has been documented, supporting our observations at the 4-month time point ^27,28^.

On the contrary, the changes in synaptic excitability point out at circuit-level changes driving the alterations in GC activity in the medium and long term. The biphasic response of GCs TBI in terms of the frequency of spontaneous excitatory postsynaptic currents aligns with previous studies and suggests an acute response that transitions into a long_-_term phase. Our results indicate that the patterning of afferent input, not merely the frequency, may contribute to the chronic hyperexcitability associated with TBI. While the loss of complexity and increased regularity might be linked to heightened synchronization and potential epileptiform activity, this hypothesis requires further investigation. In addition, niche cues such a neuroinflammation, that also shows a biphasic response ^29,30^, might play a role in TBI-associated (and sham surgery associated) changes in GC activity. In summary, our study highlights the dynamic nature of neuronal network remodelling in the DG following TBI. Understanding these temporal dynamics, and their underlying mechanisms, as well as including new parameters such as complexity, can help shape future therapeutic strategies for TBI.

## Data availability

The data underlying this article will be shared on reasonable request to the corresponding author

## Supplementary material

See supplemental information.

---

**Figure supplementary 1.**
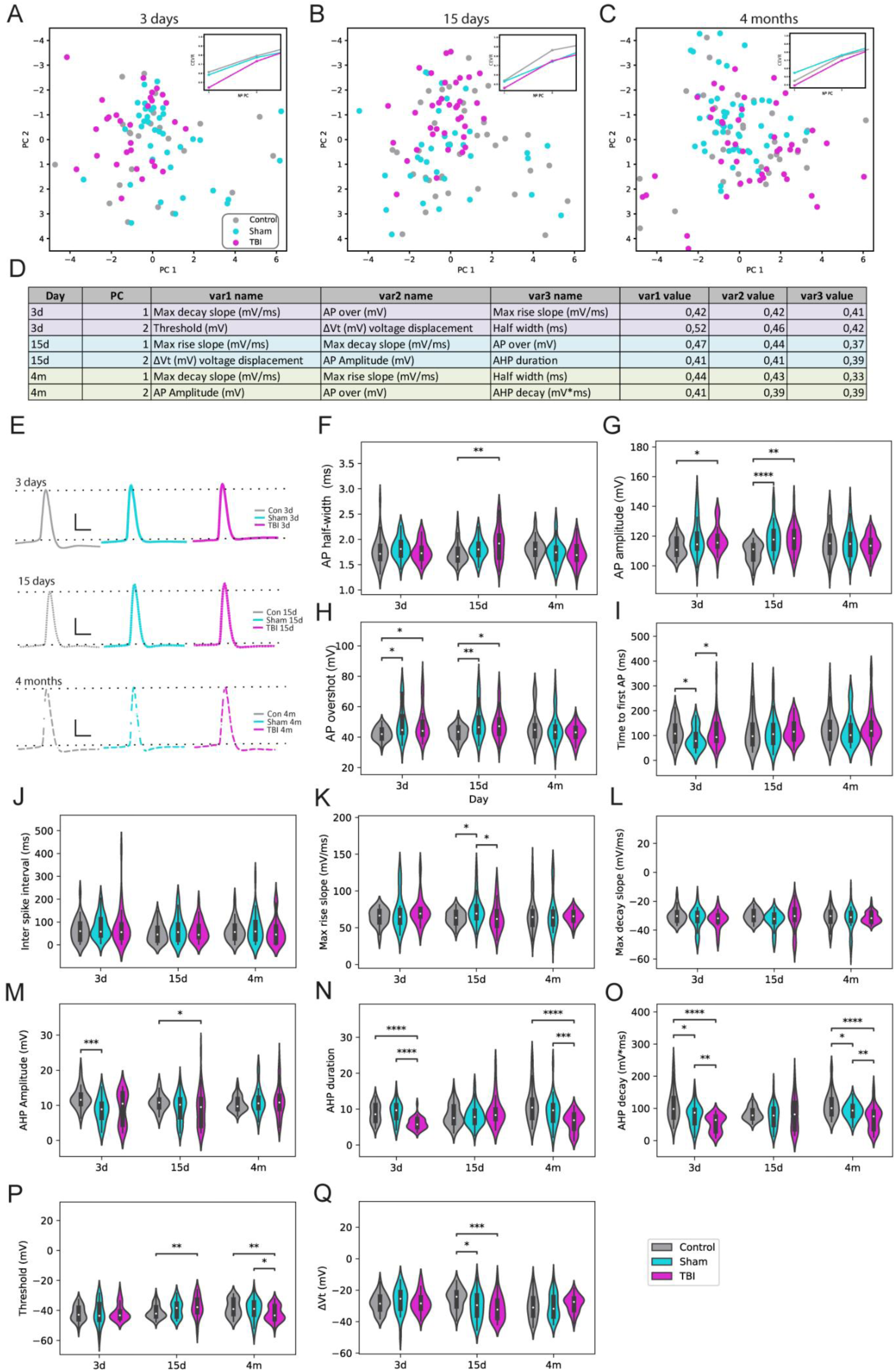
Principal Component Analysis (PCA) and Cumulative Explained Variance Ratio (CEVR) assessment of action potential features at short_-_medium- and long-term. (**A-C**) Two-dimensional PCA plots illustrating the distribution and Cumulative Explained Variance Ratio (CEVR) plots (insets) depicting the proportion of total variance explained by principal components for control, sham, and TBI granule cells at day 3 (A), day 15 (B), and 4 months (C). (**D**) Summary table presenting the loading variables with higher weights in Principal Component Analysis 1, 2, and 3 for day 3, day 15, and 4 months. (**E**) Examples of action potentials recorded 3 days, 15 days and 4 months after TBI/sham surgery with a corresponding control (Scale bars: 20 mV, 5ms). (**F**) AP half-width in control, sham and TBI groups at day3, day 15 and 4 months after the surgery (for day 15: Kruskal-Wallis test, H(2)= 9.241, p= 0.0098. ** p=0.0083 for TBI vs control in Dunn’s multiple comparisons test). (**G**) AP amplitude in control, sham and TBI groups at day 3, day 15 and 4 months after the surgery (for day 3: Kruskal-Wallis test, H(2)= 7.176, p= 0.0277. *p=0,0292 for TBI vs control in Dunn’s multiple comparisons test. For day 15: Kruskal-Wallis test, H(2)= 20.30, p <0.0001. **p=0.012 for TBI vs control and ****p<0.0001 for sham vs control in Dunn’s multiple comparisons test). (**H**) AP overshot in control, sham and TBI groups at day 3, day 15 and 4 months after the surgery (for day 3: Kruskal-Wallis test, H(2)= 9.544, p= 0.0085. *p= 0.0176 for sham vs control, ^#^p= 0,0311 for TBI vs control in Dunn’s multiple comparisons test. For day 15: Kruskal-Wallis test, H(2)= 11.36, p= 0.0034. **p=0.0051 for sham vs control and *p=0.0247 for TBI vs control in Dunn’s multiple comparisons test). **(I)** Time to first action potential (AP) in control, sham and TBI granule cells at day 3, day 15 and 4 months after the surgery (for day 3: Kruskal-Wallis test, H(2)= 9.544, p= 0.041. *p= 0.042 for control vs sham; H(2)= 21.544, p= 0.031. *p= 0.036 for sham vs TBI in Dunn’s multiple comparisons test). **(J)** Inter-spike interval in control, sham and TBI granule cells at day 3, day 15 and 4 months after the surgery. (**K**) Maximum rise slope in control, sham and TBI granule cells at day 3, day 15 and 4 months after the surgery (for day 15: Kruskal_-_Wallis test, H(2)= 12.67, p= 0.043. *p= 0.045 for control vs sham; H(2)= 11.748, p= 0.043. *p= 0.046 for sham vs TBI in Dunn’s multiple comparisons test). (**L)** Maximum decay slope in control, sham and TBI granule cells at day 3, day 15 and 4 months after the surgery. (**M**) Afterhyperpolarization (AHP) amplitude in control, sham and TBI granule cells at day 3, day 15 and 4 months after the surgery (for day 3: Kruskal-Wallis test, H(2)= 2.67, p= 0.0023, *p= 0.0025 for control vs sham. For day 15 H(2)= 6.67, p= 0.033, *p= 0.035 for control vs TBI in Dunn’s multiple comparisons test). (**N**) AHP duration in control, sham and TBI groups at day 3, day 15 and 4 months after the surgery (for day 3: Kruskal-Wallis test, H(2)= 27.06, p<0.0001. ^####^p<0.0001 for TBI vs control and ****p= 0.0001 for sham vs TBI in Dunn’s multiple comparisons test. For 4 months: Kruskal-Wallis test, H(2)= 26.76, p< 0.0001. ****p<0.0001 for TBI vs control and **p=0.0035 for TBI vs sham in Dunn’s multiple comparisons test). (**O**) AHP decay in control, sham and TBI granule cells at day 3, day 15 and 4 months after the surgery. (for day 3: Kruskal-Wallis test, H(2)= 9.536, p= 0.041. *p=0.043 for control vs sham; H(2)= 5.544, p= 0.0021. **p= 0.0026 for sham vs TBI; H(2)= 7.634, p= 0.00002. ****p= 0.000026 for control vs TBI. For month 4: H(2)= 7.365, p= 0.031. *p=0.032 for control vs sham; H(2)= 7.944, p= 0.0041. **p= 0.0046 for sham vs TBI; H(2)= 7.634, p= 0.00022. ****p= 0.000023 for control vs TBI in Dunn’s multiple comparisons test). (**P**) AP threshold in control, sham and TBI granule cells at day 3, day 15 and 4 months after the surgery. (for day 15: Kruskal-Wallis test, H(2)= 7.987, p= 0.0041. *p=0.0042 for control vs TBI. For month 4: H(2)= 6.352, p= 0.0031. **p=0.0032 for control vs TBI; H(2)= 5.782, p= 0.0021. **p= 0.0026 for sham vs TBI in Dunn’s multiple comparisons test). (**Q**) Voltage displacement in control, sham and TBI granule cells at day 3, day 15 and 4 months after the surgery. (For day 15: Kruskal-Wallis test, H(2)= 9.234, p= 0.032. *p=0.036 for control vs sham; H(2)= 7.533, p= 0.00022. ***p=0.00026 for control vs TBI in Dunn’s multiple comparisons test). n=31 neurons from control; 24 from sham and 37 from TBI at day 3; n=36 neurons from control; 36 from sham and 37 from TBI at day 15; n=49 neurons from control; 36 from Sham and 52 from TBI at 4 months. In panels F to I dots in violin plots represent individual cell values and dashed lines quartiles and medians. From panel J to Q in violin plots represent group medians and lines SD.

**Figure supplementary 2.**
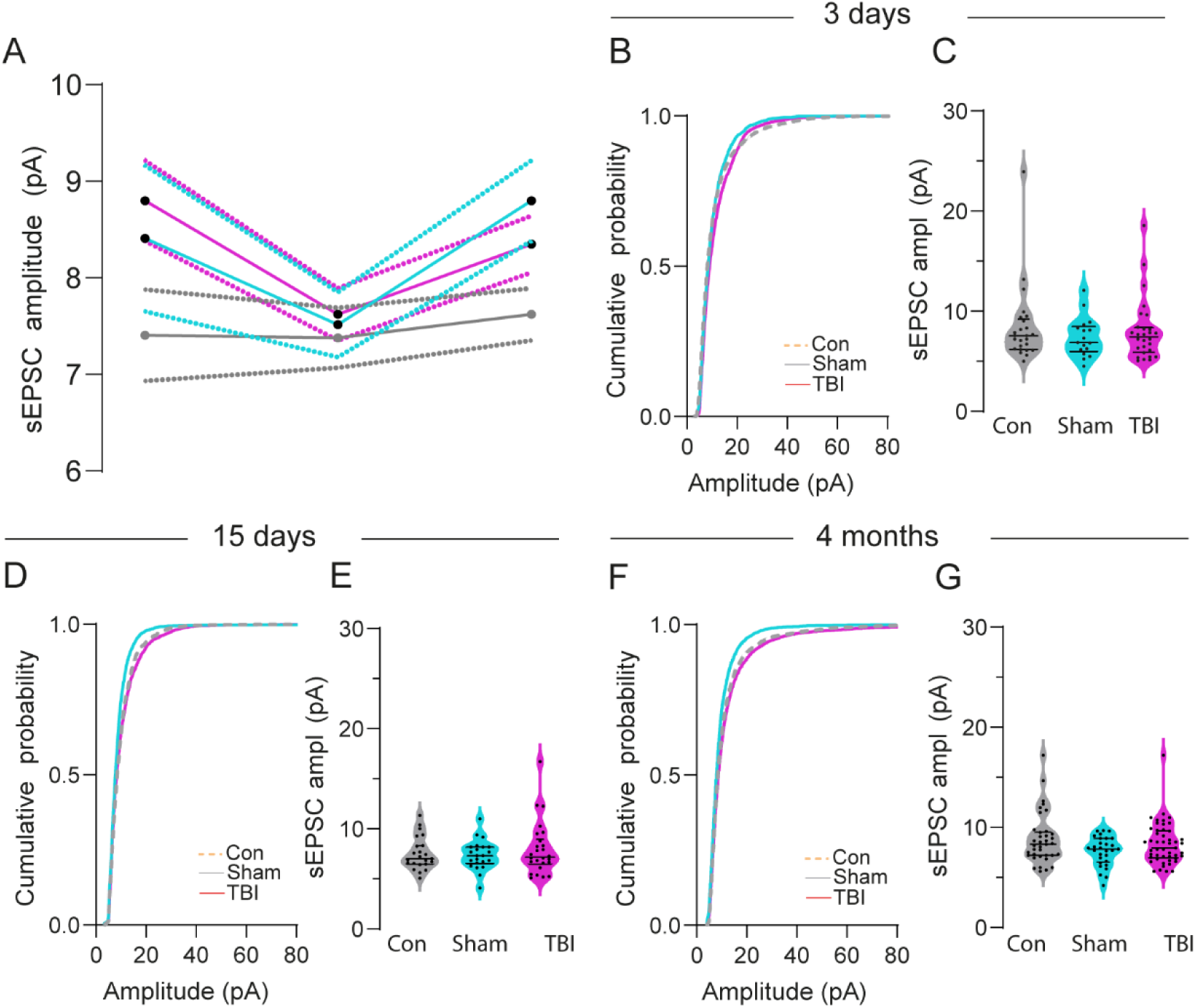
TBI does not affect amplitude of sEPSC at the short-, medium- or long-term. (**A**) sEPSC amplitude at the short_-_medium- and long-term for control, sham and TBI granule cells (**B**) sEPSC amplitude at the short_-_medium- and long_-_term for control, sham and TBI granule cells. (**B-C**) Cumulative probability plots of sEPSC amplitude (B) and violin plots showing sEPSC amplitudes (C) for control, sham and TBI granule cells at day 3 after the surgery (n=25 for control, n= 17 for sham and n= 28 for TBI). (**D-E**) Cumulative probability plots of sEPSC amplitude (D) and violin plots showing sEPSC amplitudes (E) for control, sham and TBI granule cells at day 15 after the surgery. (n=23 for control, n= 22 for sham and n= 26 for TBI. (**F-G**) Cumulative probability plots of sEPSC amplitude (F) and violin plots showing sEPSC amplitudes (G) for control, sham and TBI granule cells 4 months after the surgery (n=37 for control, n= 29 for sham and n= 49 for TBI). For line graphs dots represent group means and area fill represents SEM. Dots in violin plots represent cell median values and dashed lines quartiles and medians.

**Figure supplementary 3.**
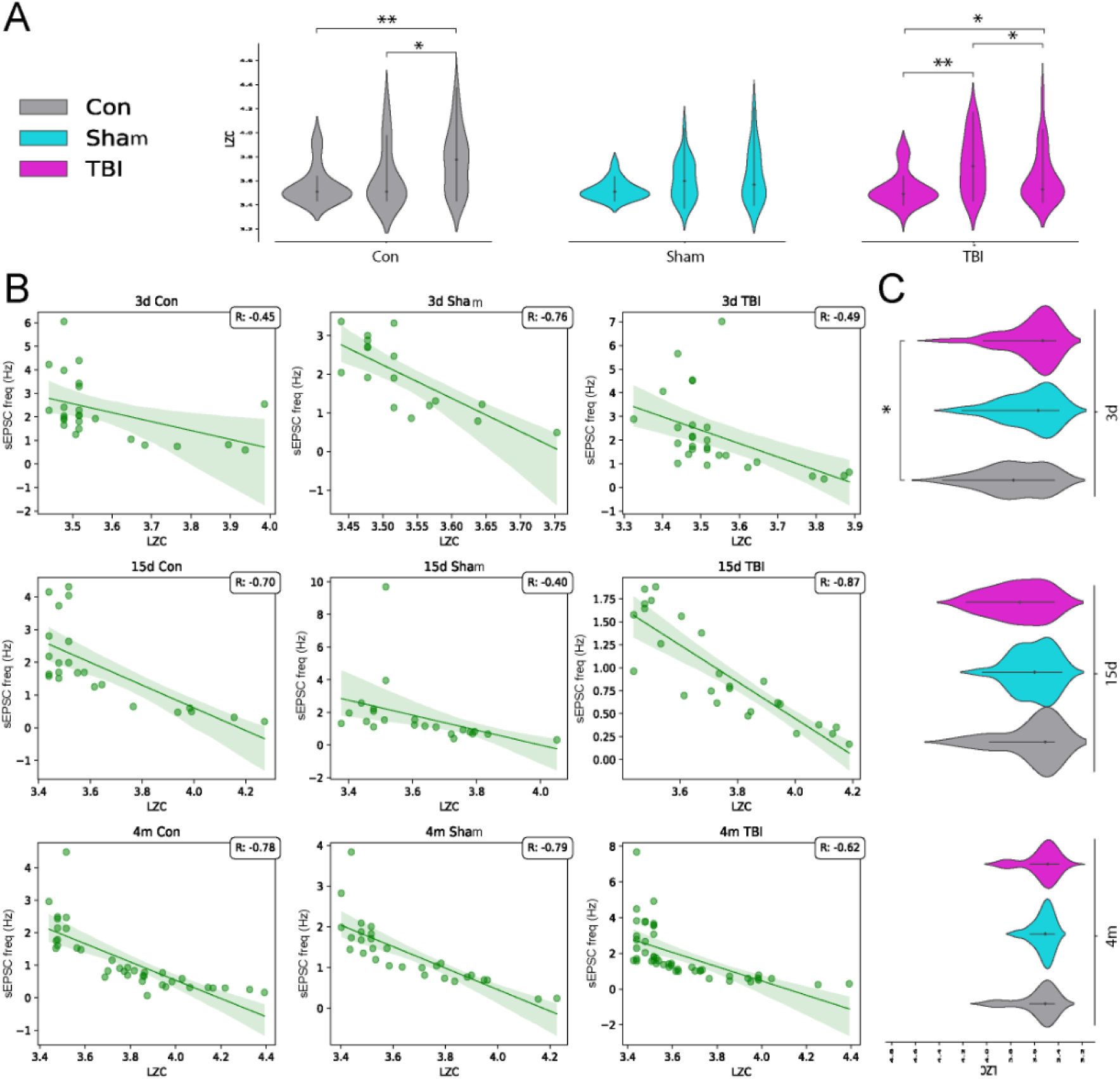
Lempel-Ziv complexity (LZC) analysis of normalized sEPSC sequences. **(A)** Comparison of LZC across different time points for control, sham, and TBI groups. (Kruskal-Wallis test, H(2)= 4.456, p=0.0254, *p= 0.0263 15 days vs 4 month in control group; H(2)= 6.678, p=0.0032, **p= 0.0031 for 3 days vs 4 month in control group; H(2)= 3.678, p=0.036. *p= 0.0412 for 15 days vs 4 month in TBI group, H(2)= 6.432, p=0.0012. **p= 0.0011 for 3 days vs 15 days in TBI group; H(2)= 4.386, p=0.0414. *p= 0.0432 for 3 days vs 4 month in TBI group in Dunn’s multiple comparisons test). **(B)** Correlation between LZC and median sEPSC over time and across groups. The correlation value (R) is indicated in the top right corner of each subplot. **(C)** Comparison of LZC between groups at specific time points. (Kruskal-Wallis test, H(2)= 5.435, p=0.0032. *p= 0.0034 for control vs TBI in 4 months in Dunn’s multiple comparisons test). n=26 neurons from control; 18 from sham and 27 from TBI at day 3; n=23 neurons from control; 22 from sham and 25 from TBI at day 15; n=36 neurons from control; 29 from sham and 48 from TBI at 4 months. Violin plots represent group medians and lines SD. In line graphs area fill represents SEM and dots represent cell values.

**Figure supplementary 4.**
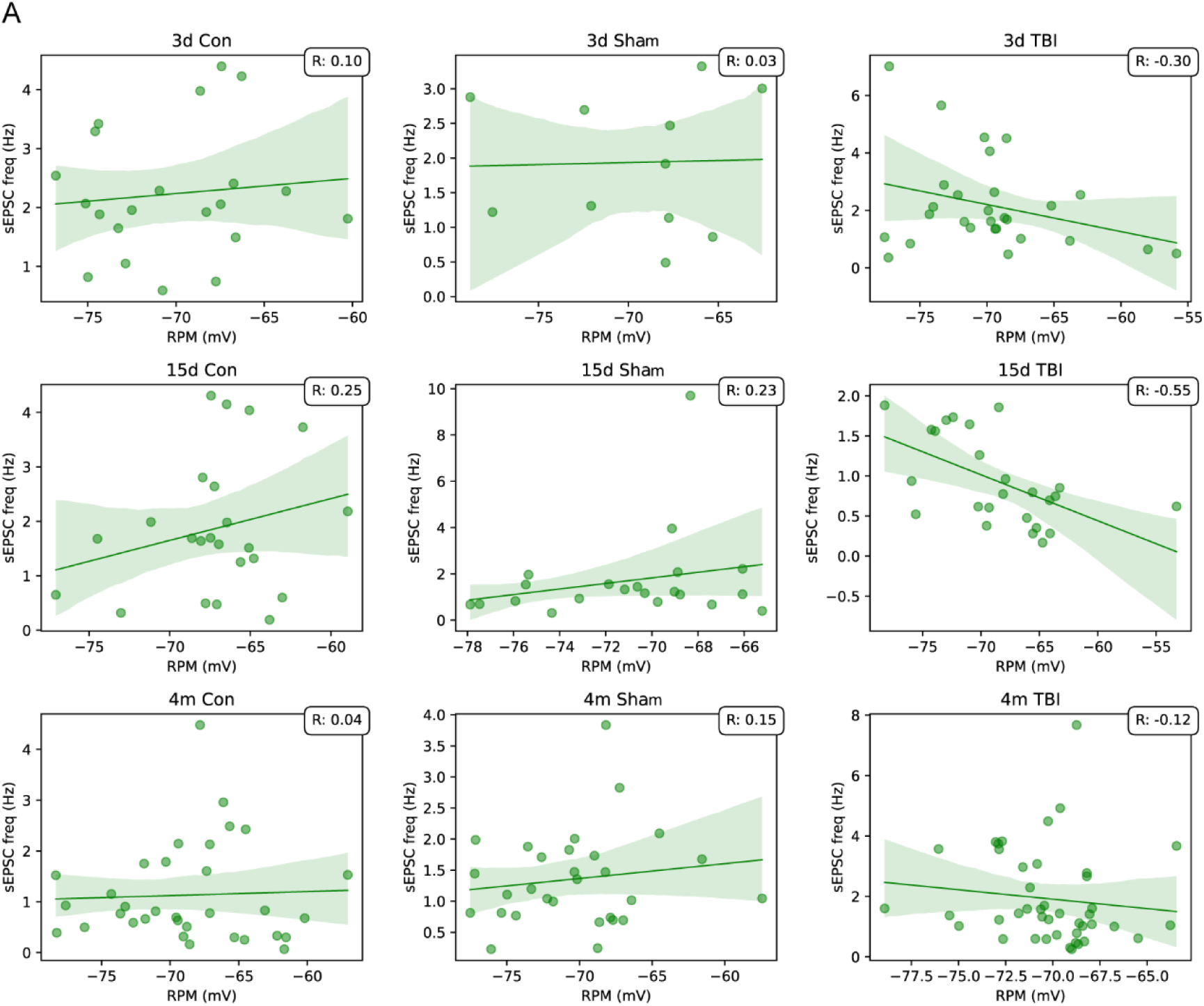
Frequency of synaptic events is not correlated with resting membrane potential of granule cells. **(A)** The correlation between sEPSC frequency and resting membrane potential is plotted for each state (left to right) and time point (top to bottom). The correlation coefficient (R) is shown in the upper right corner of each subplot. In line graphs area fill represents SEM and dots represent cell values.

