## Supplemental Files for "Biphasic Changes in Hippocampal Granule Cells after Traumatic Brain Injury"

*Supplemental material*

*Electrophysiology: whole-cell recordings*

Cell access was obtained in the voltage-clamp mode and Resting Membrane Potential (RMP) was measured immediately upon break-in in the current-clamp mode by setting the clamp current equal to zero. The firing characteristics of the recorded cells was assessed using intracellular injections of rectangular current pulses of increasing amplitude (range: - 200 pA to + 200 pA; ΔI = 20 pA; duration: 400 ms) in current-clamp mode. The first action potential (AP) that was generated with the use of this current steps protocol was used in further analysis of AP properties. For each cell, the relationship between injected current intensity and the number of AP was plotted. Evaluated AP properties were: voltage displacement from RMP to AP threshold (ΔVt), AP amplitude (AP amp), AP duration (AP dur) at 50% repolarization, AP maximum rise slope (AP max rise), AP maximum decay slope (AP max decay), afterhyperpolarization amplitude (AHP amp), and AHP duration (AHP dur) at 50% of decay back to RMP. The membrane input resistance ($R_{in}$) was determined from the slope of the current–voltage relationship. Membrane time constant *tau* was estimated by fitting the decay phase of the voltage phase to a standard exponential equation. Spontaneous excitatory postsynaptic currents (sEPSCs) were recorded as inward currents from neurons, which were voltage-clamped at -86 mV and synaptic events were recorded for 3 min for each neuron. Only cells with stable access resistance were accepted for the data analysis. Spontaneous EPSCs were detected off-line and analyzed using the Mini Analysis software (Synaptosoft, USA). Individual recorded traces were searched for peak currents using the automatic detection protocol. The average baseline current was calculated for an interval between 3 and 5 ms before the peak, the threshold amplitude for EPSC detection was set to 4 pA. Afterward recordings were inspected manually for correctness.

*Lempel Ziv Complexity*

Lempel-Ziv complexity (LZC) is a measure of the complexity of a finite sequence based on the Lempel-Ziv compression algorithm [1]. It quantifies the rate at which new patterns, or unique substrings, appear as the sequence is scanned from beginning to end. Specifically, LZC counts the number of unique substrings encountered during this process, reflecting the repetitiveness and structure of the sequence.

To calculate LZC, the sequence is parsed sequentially and each new unique substring is identified and added to a dictionary of known substrings. The complexity is then given by the total number of unique substrings detected. A higher LZC indicates a more complex sequence with less redundancy, while a lower LZC indicates greater regularity and predictability.

To estimate the complexity of a time series X= {$x_t$; t=1, ... , T}, sequence *X* is parsed into a number *W* of words by considering any subsequence that has not yet been encountered as a new word. The Lempel-Ziv complexity, represents the smallest number of distinct words *W*, needed to fully reconstruct the information in the original time series. For example, the sequence *1001111011000010* can be parsed in 6 words *1, 0, 01, 1110, 1100, 0010* giving a complexity LZC=6. An easy way to apply the Lempel-Ziv algorithm can be found in [2]. The LZC can be normalized based in the length T of the discrete sequence and the alphabet length $\alpha$ as $\mathrm{LZCn} = \frac{LZC\,[\log_{\alpha} T]}{T}$.

*Principal component Analysis (PCA)*

PCA, a widely used dimensionality reduction technique in data analysis, is a powerful tool for understanding and simplifying complex datasets with many features, as in this case. It provides a concise representation of the data while preserving maximum information. PCA identifies principal components (PCs), which are linear combinations of the original features. Each principal component captures a certain amount of variance in the data. Typically, the original principal components capture the majority of the variance, facilitating dimensionality reduction [3].

However, when applying PCA, it is crucial to determine how much of the variation in the data is explained by each principal component. A commonly used tool to achieve this is the cumulative explained variance ratio (CEVR). This metric is the cumulative sum of the explained variance of each principal component included in the analysis in turn. The CEVR proves to be instrumental in discerning the total variance retained as additional principal components are included. It helps to capture the trade-off between dimensionality reduction achieved by using fewer components and the retention of information from the original dataset.

Furthermore, we are interested in identifying the variables that contribute most to the components in the PCA analysis. For this purpose, we extract the principal loadings, which are essentially the top eigenvectors of the covariance matrix in the PCA algorithm.

In our study, we used PCA to reduce the dimensionality of the entire data set, focusing on passive membrane properties. Two principal components were obtained to facilitate comparisons between three different time points. In addition, we examined the three highest loadings to identify the original measures that were most influential in this comparative analysis.
